# Year-round dynamics of amplicon sequence variant communities differ among eukaryotes, *Mimiviridae*, and prokaryotes in a coastal ecosystem

**DOI:** 10.1101/2021.02.02.429489

**Authors:** Florian Prodinger, Hisashi Endo, Yoshihito Takano, Yanze Li, Kento Tominaga, Tatsuhiro Isozaki, Romain Blanc-Mathieu, Yasuhiro Gotoh, Hayashi Tetsuya, Etsunori Taniguchi, Keizo Nagasaki, Takashi Yoshida, Hiroyuki Ogata

## Abstract

Coastal seawater is the habitat of different microbial communities. These communities are affected by seasonal environmental changes and fluctuating nutrient availability, as well as competitive and cooperative interspecific interactions. In this work, we investigated the seasonal dynamics of communities of eukaryotes, a major group of double-stranded DNA viruses infecting eukaryotes (i.e. *Mimiviridae),* as well as prokaryotes in the Uranouchi Inlet, Kochi, Japan. We performed metabarcoding using ribosomal RNA genes and the *Mimiviridae polB* gene as marker genes in 43 seawater samples collected during 20 months. Communities characterized by the compositions of amplicon sequence variants (ASVs) showed synchronic seasonal cycles for eukaryotes, *Mimiviridae,* and prokaryotes. However, the community dynamics showed intriguing differences in several aspects such as the recovery rate after a year. We further show that the differences in the community dynamics can be explained by differences in the recurrence/persistence levels of individual ASVs among eukaryotes, *Mimiviridae,* and prokaryotes. *Mimiviridae* ASVs were less persistent than eukaryotic ASVs, and prokaryotic ASVs were the most persistent. We argue that the differences in the specificity of interactions (i.e. virus-eukaryote *vs* prokaryote-eukaryote) as well as the survival strategies are at the origin of the distinct community dynamics among eukaryotes, *Mimiviridae,* and prokaryotes.

**One sentence summary:** A one year observation of coastal microbial communities revealed similar but different community dynamics for eukaryotes, a group of large viruses, and prokaryotes.

## Introduction

Microbes – cellular microorganisms and viruses – in marine ecosystems are responsible for biogeochemical cycling (Not *et al*. 2012; Brum and Sullivan 2015; Fuhrman, Cram and Needham 2015), which is driven by diverse members of microbial communities. The compositions of microbial communities change over time and space in response to environmental factors and through species interactions. Therefore, to understand a marine ecosystem, it is fundamentally important to study the structures of microbial communities, their dynamics, and the mechanism that drives the change of microbial communities in the ecosystem.

Previous studies have explored the temporal dynamics of eukaryotic (Chen *et al*. 2017; Giner *et al*. 2019; Gran-Stadniczeñko *et al*. 2019a), prokaryotic (Gilbert *et al*. 2012; Fuhrman, Cram and Needham 2015; Milici *et al*. 2016; Sakami *et al*. 2016; Teeling *et al*. 2016; Ward *et al*. 2017), both eukaryotic and prokaryotic (Bock *et al*. 2018; Chafee *et al*. 2018), and viral communities (Ignacio-Espinoza, Ahlgren and Fuhrman 2020). Other studies have investigated co-variations among these cellular communities (Needham and Fuhrman 2016; Martin-Platero *et al*. 2018; Santi *et al*. 2019) or cellular and viral communities (Chow and Fuhrman 2012; Needham *et al*. 2013; Pagarete *et al*. 2013; Johannessen *et al*. 2017; Arkhipova *et al*. 2018; Sandaa *et al*. 2018; Gran-Stadniczeñko *et al*. 2019b). Overall, these studies have revealed similarities and differences in community dynamics among different microbe groups. Yearly cycles and the recurrence of certain operational taxonomy units at 12-month intervals have been detected for eukaryotes in the north western Mediterranean Sea (Giner *et al*. 2019), prokaryotes in the English Channel (Gilbert *et al*. 2012; Fuhrman, Cram and Needham 2015), and small viruses at a near-coastal site of southern California (Ignacio-Espinoza, Ahlgren and Fuhrman 2020) through long-term observations. A short-term study that compared prokaryotic and eukaryotic communities in Massachusetts Bay revealed a higher rate of decline in community similarity for eukaryotes than for prokaryotes (Martin-Platero *et al*. 2018). Furthermore, changes in abiotic factors were found to have a lower impact on the eukaryotic community than on prokaryotic communities in the Mediterranean Sea (Giner *et al*. 2019).

In this study, we investigated the community structures of microbial eukaryotes, viruses in the family *Mimiviridae* (phylum *Nucleocytoviricota),* and prokaryotes in the Uranouchi Inlet in Kochi prefecture, Japan. The Uranouchi Inlet is a semi-enclosed, eutrophic inlet, which is located at the south eastern side of Shikoku Island, Japan. The climate of Shikoku Island is humid subtropical and shows clear seasonality, with contrasting high and low water temperatures in summer and winter, respectively. The contrasting seasons provide abiotic conditions that change throughout the year. The inlet is known for recurrent harmful blooms of eukaryotic microalgae including *Karenia mikimotoi* (Dinophyceae), *Chattonella* spp. (Raphidophyceae), and *Heterosigma akashiwo* (Raphidophyceae), which occasionally damage the local fishery and aquaculture. No previous studies have investigated the dynamics of the microbial eukaryotic community in this inlet.

Viruses are thought to be important top-down regulators of communities of microorganisms, including eukaryotes. For example, the collapse of eukaryotic phytoplankton blooms is associated with viruses (Tarutani, Nagasaki and Yamaguchi 2000; Tomaru *et al*. 2004; Lehahn *et al*. 2014; Moniruzzaman *et al*. 2014). The sudden increase of virions during the period of bloom decline suggests that viruses are top-down regulators of phytoplankton populations (Nagasaki *et al*. 2003; Kuhlisch *et al*. 2020). The family *Mimiviridae* is a major group of large DNA viruses that infect diverse eukaryotes in marine environments (Hingamp *et al*. 2013). A recent study revealed an association between eukaryote and *Mimiviridae* communities in a global metagenomic dataset (Endo *et al*. 2020). Another study detected cooccurring pairs of eukaryotes and large DNA viruses including members of the *Mimiviridae* through time-series sampling for more than one year in Northern Skagerrak, South Norway (Gran-Stadniczeñko *et al*. 2019b). These spatial and temporal associations suggest the tight ecological interactions between diverse *Mimiviridae* and eukaryotes, that is, the obvious dependence of *Mimiviridae* replication on the activity of their hosts and the possible top-down regulatory roles of these viruses on the dynamics of the eukaryote community as a consequence of lytic infection. However, long-term studies on the associations between these viruses and eukaryotes are still limited. Thus, how their community dynamics are related or distinct is under-investigated. We recently established a highly sensitive metabarcoding method for *Mimiviridae,* which successfully detected diverse *Mimiviridae* in the Uranouchi Inlet (Prodinger *et al*. 2020). In this study, the community of *Mimiviridae* was analyzed with this method for the first time on a time-series sample set and compared with communities of microbial eukaryotes to see if they show synchronic seasonal dynamics in the study area towards better understanding their ecological interactions.

We also investigated the dynamics of the prokaryotic community. *Mimiviridae* and eukaryotes directly interact with each other through infection, while eukaryotes and prokaryotes interact with each other either directly by grazing and parasitism, or indirectly by metabolization of cell debris and nutrient-rich cellular exudates. An example of the indirect interaction between eukaryotes and prokaryotes can be seen during algal blooms. The decline of algal blooms is associated with the release of cellular debris and other organic substrates (Kuhlisch *et al*. 2020), which become important resources for heterotrophic prokaryotes (Gómez□Pereira *et al*. 2012). Therefore, we reasoned that the prokaryotic community serves as a contrasting reference for the comparison of communities between directly interacting eukaryotes and *Mimiviridae.*

We performed metabarcoding of eukaryotes, prokaryotes, and *Mimiviridae* using 18S and 16S rRNA genes and the *Mimiviridae polB* gene as marker genes, based on 43 seawater samples collected during 20 months from January 2017 to September 2018. Microbial communities were characterized at the amplicon sequence variant (ASV) level, which is the highest possible resolution for operational taxonomic grouping of amplicon sequences. We characterized the similarities and differences in community dynamics among eukaryotes, *Mimiviridae*, and prokaryotes.

## Materials and Methods

### Sampling and DNA extraction

Seawater was collected at four stations in Uranouchi Inlet, Kochi Prefecture, Japan (i.e. station “j”: 33°25’43.2”N 133°22’49.5”E, station “m”: 33°25’60.0”N 133°24’38.3”E, station “f”: 33°26’33.6”N 133°24’41.8”E, station “u”: 33°25’49.7”N 133°24’01.4”E). Time-series sampling was performed sporadically (average interval, 15 days) with a higher sampling frequency during summer. In total, 43 sample sets (i.e. 43 seawater samples, with three size fractions each) were obtained over 20 months from 5^th^ January 2017 (170105-j) to 25^th^ September 2018 (180925-j). The samples were denoted with the sampling date followed by the station’s abbreviation in “yymmdd-station” format.

Seawater (10 L) was collected from 5-m depth, transported to the laboratory, and then filtered through 3.0-μm (diameter 142 mm) (polycarbonate, Merck, Darmstadt, Germany) and 0.8-μm (diameter 142 mm) membranes (polycarbonate, Merck). The filtrate (1 L) was further filtered through a 0.22-μm filtration unit (Sterivex, polycarbonate, Merck). The filters were kept at −80°C until DNA extraction with the Proteinase-K method for the 0.22-μm filtration units (Frias-Lopez *et al*. 2008) and the xanthogenate-sodium dodecyl sulfate method for larger size fractions (0.8-μm and 3-μm filters) (Yoshida *et al*. 2003).

### PCR amplification

The 18S ribosomal RNA gene primers (forward “V8f”: ATAACAGGTCTGTGATGCCCT, reverse “1510r”: CCTTCYGCAGGTTCACCTAC) (Bradley, Pinto and Guest 2016) targeting the V8/V9 region were used on both the 0.8–3 μm and >3 μm size fractions. Hereafter, the 0.8–3 μm and >3 μm fractions are referred to as the PicoNano and NanoPlus fractions, respectively. The PCR analysis was performed by mixing 5 μL of 1 μmol·L^-1^ of both primers and 2.5 μL of 0.25 ng·μL^-1^ template DNA with 12.5 μL 2× KAPA HiFi HotStart ReadyMix (Roche, Basel, Switzerland). The thermal program was 98°C for 3 min, followed by 25 cycles of 98°C for 20 s, 65°C for 15 s, 72°C for 15 s, and 72°C for 10 min. The 16S rRNA gene amplification was performed by mixing 5 μL of 1 μmol·L^-1^ of both primers (forward “Pro-16S NGS”: CCTACGGGNBGCASCAG, reverse “Pro-16S NGS”: GACTACNVGGGTATCTAATCC) (Takahashi *et al*. 2014) with 2 μL of 1 ng·μL^1^ sample DNA (0.2-0.8 μm size fraction) and 12 μL 2× KAPA HiFi HotStart ReadyMix. The thermal program was 95°C for 3 min, then 25 cycles of 95°C for 30 s, 55°C for 30 s, 72°C for 30 s, and final extension at 72°C for 5 min. For the *Mimiviridae* DNA polymerase gene amplification, we used a recently published method that has been described in detail elsewhere (i.e. “MP10v2” in combination with “protocol 3”) (Prodinger *et al*. 2020).

### Clean up and sequencing

The 18S rRNA gene and 16S rRNA gene amplicons were purified with Agencourt AMPure XP beads (Beckman Coulter, Inc., Brea, CA, USA) following Illumina’s library preparation protocol. The *Mimiviridae polB* gene amplicons were purified by gel extraction (Prodinger *et al*. 2020). After amplicon purification, dual indices were attached by PCR. The second purification was performed according to Illumina’s library preparation protocol for the cellular amplicons and with gel extraction for *polB* amplicons. The final library concentration was 10 pM with a 30% PhiX spike. The sequencing data were generated with paired-end sequencing (2 × 300 nucleotides) on the MiSeq platform (Illumina, Inc., San Diego, CA, USA).

### Analysis of prokaryotic and eukaryotic amplicon raw reads

The data were analyzed separately using QIIME 2 (Bolyen *et al*. 2018) with default settings unless noted otherwise. The pipelines for both prokaryotes and eukaryotes were nearly identical. The raw reads were imported to QIIME 2 (version 2020.2.0). dada2 (Callahan *et al*. 2016: 2) was used for ASV generation in QIIME 2, mostly with default settings. This included several steps. Primers were removed according to length and reads were corrected according to the dada2 algorithm. The final low quality basepairs of each read were removed (truncation at 240 bp for18S data and at 250bp for 16S data). Finally, dada2 merged forward and reverse reads and removed chimeric sequences. The ASVs were taxonomically assigned at 90% nucleotide identity using “classify-consensus-vsearch” in QIIME 2 against the SILVA 132 database (taxonomic majority, 97%) for either 16S amplicon reads or 18S amplicon reads.

### Analysis of *MimiviridaepolB* amplicon raw reads

*Mimiviridae polB* ASVs were generated by a newly developed pipeline named MAPS2, which is mostly based on R (3.6.3) (www.R-project.org, 17 February 2021, date last accessed) and the “dada2” package. The functions of dada2 were used with default settings unless specified otherwise. Raw reads were loaded using the “filterAndTrim” command, truncated at a length of 240 bp (both forward and reverse reads), and 2 expected read errors were accepted. Primer sequence regions were trimmed (i.e. forward reads: 40 bp and reverse reads: 38 bp). The chance of sequencing error was calculated with dada2’s “learnErrors” function and sequences were dereplicated with the “derepFastq” function. The “dada” function was used on the reads, and the reads were merged with the “mergePairs” function. Chimeric sequences were removed using the “removeBimeraDenovo” function. The nucleotide sequences and the ASV table were then exported. The nucleotide sequences were searched against a database of 11,413 *polB* sequences, 1,004 of which were viral *Mimiviridae* sequences, using BLASTx (2.9.0)(Altschul *et al*. 1990) (like in the original MAPS (Li *et al*. 2018)). Sequences with the best hit to viral sequences were processed further (E-value <10^-5^). Both the original nucleotide sequence and the amino acid sequences (translated by BLASTx) were saved. MAFFT (7.453) (Katoh and Standley 2013) was used to align the reads with those in a reference file containing 1109 PolB sequences from different organisms and viruses. The aligned PolB sequences were added to a reference tree (Endo *et al*. 2020) using pplacer (1.1.alpha19) (Matsen, Kodner and Armbrust 2010). We removed ASVs that were not within the *Mimiviridae* branch of the reference tree with R (Fig. S5). The ASVs within the *Mimiviridae* branch were split into 13 clades for community structure analysis (Fig. S5). Finally, a Python (3.7.5) (www.python.org, 17 February 2021, date last accessed) script was used to cut the *Mimiviridae* ASV sequences in a common region. The ASVs that became identical as a result of this process were clustered using cd-hit (clustering at 100% nucleotide identity).

### Community composition and diversity analysis

Communities were compared with R with the “vegan” (2.5-6) (Oksanen *et al*. 2012) and “ggplot2” (3.2.1) (https://ggplot2.tidyverse.org, 17 February 2021, date last accessed) packages. Singleton ASVs and ASVs that were not taxonomically assigned and datasets with fewer than 8000 taxonomically assigned reads were removed before analyses of community composition (Fig. 1). Ecological analysis was performed after subsampling with “rarefy” at 8000 reads. An ASV was determined to be present in the dataset if it yielded more than 0.1% of reads after subsampling.

**Figure 1:**
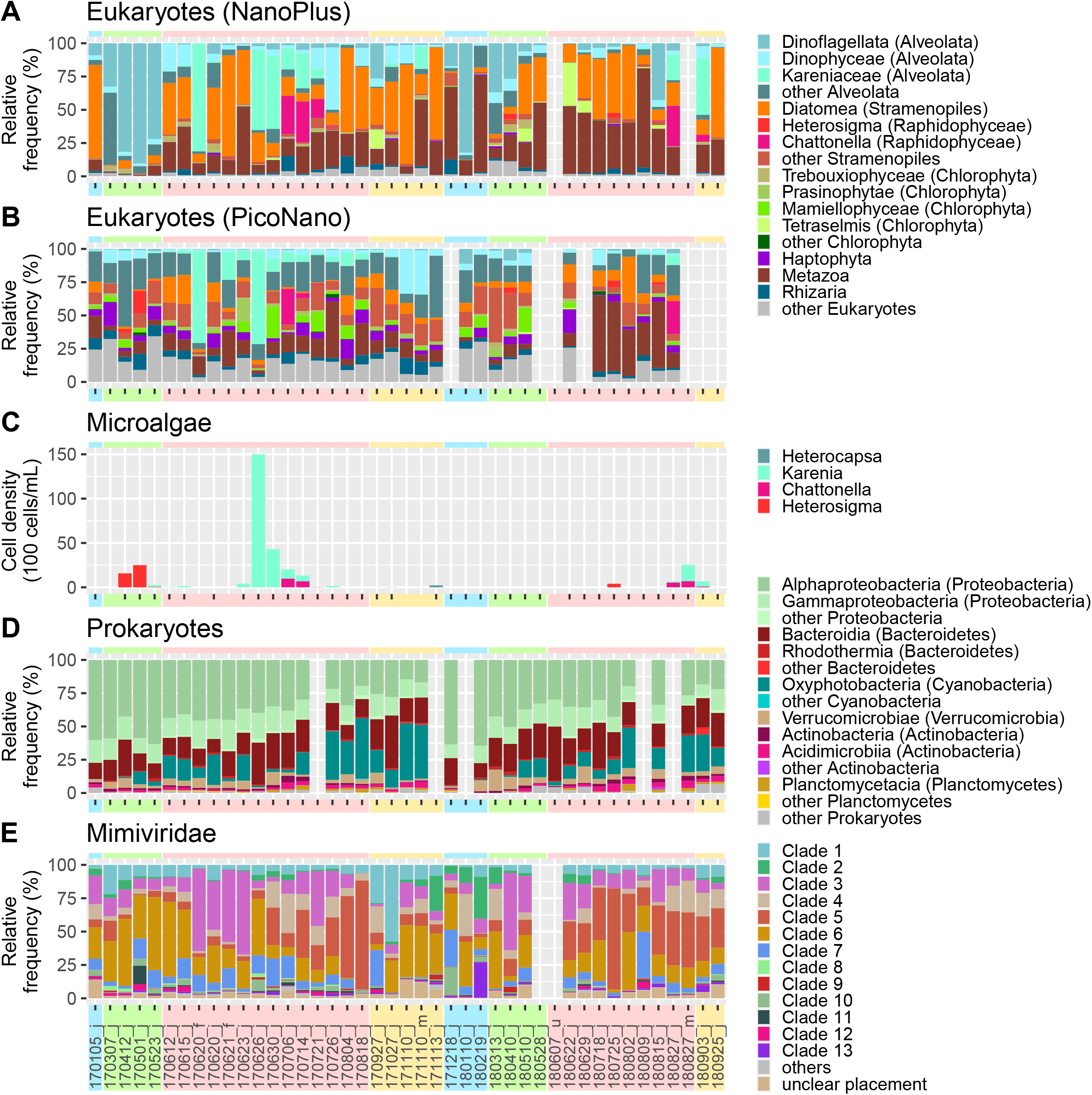
Relative abundance of different eukaryotic lineages, prokaryotic phyla, and viral clades. Bar plots show relative read percentages of generated sequencing data for eukaryotic lineages of (A) NanoPlus and (B) PicoNano size fraction, and (C) daily cell count of HAB species. (D) Relative abundance of prokaryotic phyla. (E) Relative abundance of *Mimiviridae.* Colors in the background represent seasons (blue: winter, green: spring, red: summer, and yellow: fall).

Community dissimilarity was measured using the Sørensen-Dice dissimilarity test with vegan’s “vegdist” function (method = “bray”, binary = T), as follows:

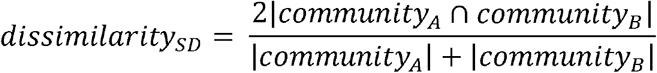

Mantel test was performed using R’s “mantel” command (5 million permutations). The NMDS analysis was conducted with R’s “cmdscale” command, and stress was calculated using the “sammon” command (MASS package). The similarity of communities among seasons was calculated by binning datasets of several months: winter (December to February), spring (March to May), summer (June to August), and fall (September to November). The *p* values for seasonal differences were calculated with R’s “t.test” command, if more than 100 values were compared we used the “pairwise.t.test” command with Benjamini–Hochberg correction.

### Cell counts, biotic, and abiotic data

Environmental metadata (temperature, salinity, phytoplankton cell counts, and concentrations of nutrients, organic and inorganic forms of nitrogen and phosphorus, dissolved oxygen, and chlorophyll *a)* for the Uranouchi Inlet from 2013 to 2019 were obtained from the observation records of the Kochi Prefectural Fisheries Research Institute with appropriate ethics approval. Since these records aim to monitor and forecast HABs in the sampling area, cell count data were only collected for HAB species. We used the daily maximum cell count data in this study. Different species are considered to be blooming at different cellular concentrations, because bloom identification is linked to heightened fish mortality according to the Prefectural Fisheries Research Institute (https://www.pref.kochi.lg.jp/soshiki/040409, 17 February 2021, date last accessed). Fish mortality rates increase when *Chattonella* spp. reaches cellular concentrations as low as 10 to 100 cells mL^-1^; when *K. mikimotoi* (an unarmored, solitary dinoflagellate, 24–40 μm long, 20–32 μm wide (Omura *et al.))* reaches concentrations ranging from several 100 to 1000 cellsomL^-1^; and when *H. akashiwo* (kidney shaped, motile dinoflagellate, 8-25 μm long (Omura *et al.))* reaches concentrations starting from 10.000 cells·mL^-1^.

## Results

### Seasonal variations in abiotic and biotic environmental factors

Abiotic factors (i.e. temperature and salinity) and the concentrations of nutrients (i.e. nitrogen species and phosphate), dissolved oxygen, and chlorophyll *a* were recorded over 6 years (Fig. S1). Temperature and salinity showed the most pronounced seasonality among all recorded biotic and abiotic factors. The differences in seasonality were less pronounced for other factors, although the concentrations of nutrients and chlorophyll *a* were higher in summer and autumn than in winter and spring (Fig. S2).

### Seasonal variation of bloom-forming phytoplankton

The abundance of six harmful algal bloom (HAB) -forming phytoplankton (i.e. *Heterocapsa circularisquama* (Dinophyceae), *K. mikimotoi* (Dinophyceae), *Cochlodinium polykrycoides* (Dinophyceae), *Chattonella* spp. (Raphidophyceae), *H. akashiwo* (Raphidophyceae), and *Pseudochattonella verruculosa* (Ochrophyta, Dictyochophyceae)) were recorded by microscopic cell counting from 2013 to 2019. Blooms of three species (i.e. *H. akashiwo, K. mikimotoi, Chattonella* spp.) occurred in the inlet and showed an approximately seasonal pattern: *H. akashiwo* bloomed in late spring (April or May), *K. mikimotoi* bloomed in early summer (~June), and *Chattonella* spp. bloomed in the middle of summer (~July). This pattern was observed in 5 of the 6 years we monitored (Fig. S2M). In 2014, 2015, and 2018, a second *H. akashiwo* bloom occurred (Fig. S1G). The highest daily cell counts of these species were in 2018 or 2019 (Fig. S1G). Blooms of *H. circularisquama, C. polykrycoides,* and *P. verruculosa* did not occur every year and their maximum abundances were relatively low (29–4900 cells·mL^-1^) (Fig. S1G, and Fig. S2M).

### Overview of sequence data

We generated from 36 to 43 metabarcoding datasets for four different communities (i.e. eukaryotic NanoPlus (>3 μm) and PicoNano (0.8–3 μm), prokaryotes (0.2–0.8 μm), and *Mimiviridae* (0.2–0.8 μm)) from a total of 43 sample sets. After discarding datasets with <8000 reads, 36 to 41 datasets were retained for each community (i.e. eukaryotic NanoPlus community: 41; eukaryotic PicoNano community: 36; prokaryotes: 38; *Mimiviridae:* 41). These metabarcoding datasets were composed of 5.2 million eukaryotic reads, 3.4 million prokaryotic reads, and 5.7 million *Mimiviridae* reads (Table 1). The subsampled datasets showed similar curve shapes and slopes at 8,000 reads, indicating sufficient sequencing depth for all four communities (Fig. S3).

**Table 1:** Summary of generated datasets. The table contains read and ASV numbers for all datasets with >8000 reads. Singletons and unassigned ASVs were removed from datasets. Average richness is shown with one standard deviation.

|  | Raw reads | Reads | Richness<br>all reads | Richness<br>per samples<br>subsampled | Average<br>richness<br>subsampled |
| --- | --- | --- | --- | --- | --- |
| **NanoPlus** | 3 613 399 | 3 001 267 | 3356 | 52 – 253 | 120 ± 51 |
| **PicoNano** | 2 692 119 | 2 180 138 | 4559 | 115 – 410 | 278 ± 68 |
| **Prokaryotes** | 4 044 427 | 3 427 158 | 4478 | 141 – 415 | 275 ± 57 |
| **Mimiviridae** | 45 106 746 | 5 712 580 | 6261 | 108 – 394 | 246 ± 72 |

The eukaryotic reads were grouped into 6413 ASVs. Only 1501 ASVs (23.4%) were represented in both NanoPlus (3356 ASVs) and PicoNano (4559 ASVs) datasets, but those ASVs accounted for 97.3% and 90.1% of total reads in the NanoPlus and PicoNano datasets, respectively. The *Mimiviridae* reads were grouped into 6261 ASVs, whilst the prokaryotic reads were grouped into 4478 ASVs. The subsampled datasets also showed relatively low richness for eukaryotes in the NanoPlus size fraction (1486 ASVs) compared with eukaryotes in the PicoNano size fraction (2184 ASVs), prokaryotes (2383 ASVs) and *Mimiviridae* (3693 ASVs). Shannon’s diversity fluctuated throughout the sampling period (Fig. S4). The eukaryotic community in the NanoPlus size fraction was less diverse than other communities during most of the sampling period (Table 1, Fig. S4).

### Composition of eukaryote, prokaryote, and *Mimiviridae* communities

Eukaryotic ASVs were classified into 16 groups (from phylum to genus levels) as shown in Figure 1. These 16 groups accounted for 98.7% of the eukaryotic reads from the NanoPlus dataset (Fig. 1A). The group with the highest relative read abundance was Metazoa (28.6%, 625 ASVs), followed by Diatomea (25.1%, 234 ASVs) and Dinoflagellata (22.5% including Dinophyceae, 469 ASVs). These three groups accounted for 77.0% of total reads. Each of the other 13 groups accounted for 0.06% to 6.6% of total reads. The relative abundance of ASVs of HAB-forming genera and families, namely *Heterosigma, Chattonella,* and Kareniaceae, was 0.26%, 2.1%, and 5.8%, respectively (Fig. 1A and Fig. 1B). Each of these groups was dominated by a single ASV; the most abundant ASV in each group accounted for more than 96.1% of the total reads in that group.

The 16 selected eukaryotic groups accounted for 88.8% of the eukaryotic reads in the PicoNano dataset. The most abundant groups were “other Alveolata” (Alveolata that were not classified as Dinoflagellata; 17.5%, 1201 ASVs), Metazoa (17.2%, 376 ASVs), and “other Stramenopiles” (Stramenopiles that were not classified as Diatomea or Raphidophyceae, 12.4%, 716 ASVs); these three groups accounted for 47.1% of all reads (Fig. 1B). Each of the remaining 13 groups contributed from 0.14% to 7.7% of total reads. The HAB-forming groups Kareniaceae, *Chattonella,* and *Heterosigma* accounted for 7.7%, 1.5%, and 0.75% of total reads, respectively. The most abundant ASVs in these three groups contributed more than 88% of their respective groups. Diatomea, Dinophyceae, and Haptophyta contributed 7.3%, 6.4%, and 5.9% of total reads, respectively. All other groups contributed less than 5%. Chlorophytes (including Trebouxiophyceae, Prasinophytae, Mamiellophyceae, and Tetraselmis) accounted for 7.2% of all reads.

Prokaryotic reads were dominated by three phyla (i.e. Proteobacteria, Bacteroidetes, and Cyanobacteria), which represented 89.3% of all prokaryotic reads (Fig. 1D). The most abundant phylum was Proteobacteria (2164 ASVs and 53.6% of total reads), 72.1% of which belonged to Alphaproteobacteria. Bacteroidetes (992 ASVs and 20.4% of total reads) was dominated by Bacteroidia (967 ASVs, 91.4% of Bacteroidetes reads). Cyanobacteria was dominated by Oxyphotobacteria (99.9% of Cyanobacteria reads). Three other phyla (Verrucomicrobiae, Actinobacteria, and Planctomycetes) contributed 4.8%, 3.8%, and 0.5% of total prokaryotic reads, respectively. Overall, these six phyla contributed 98.4% of prokaryotic reads.

We classified the 5701 *Mimiviridae* ASVs (of the 6261 total *Mimiviridae* ASVs) into 13 clades based on their phylogeny (Fig. S5). The majority of amplicon reads (61.9%) were assigned to clades 6 (518 ASVs, 24.0%), 3 (665 ASVs, 20.7%), and 5 (535 ASVs, 17.2%). The other ten clades contributed from 0.3% to 8.4% of total *Mimiviridae* reads. Six out of the 13 *Mimiviridae* clades included previously studied *Mimiviridae* (clades 1, 3, 8, 9, 10, 13) (Fig. S5). Of these six clades, clade 3, which includes Organic lake phycodnavirus 1 and 2 (Zhang *et al*. 2015), had the largest number of reads (20.7%). Clade 1 (7.8% of *Mimiviridae* reads) included haptophytes (Prymnesiophyceae) infecting *Mimiviridae* (Santini *et al*. 2013; Gallot-Lavallée *et al*. 2015; Johannessen *et al*. 2015). Clade 10 (2.4% of *Mimiviridae* reads) contained the predatory Choanoflagellate infecting virus ChoanoV1 (Needham *et al*. 2019). Each of the other clades (clades 8, 9, 13) with known *Mimiviridae* contributed less than 2% of *Mimiviridae* reads.

### Synchronic seasonal changes in microbial communities

The non-metric multidimensional scaling (NMDS) ordination of the datasets revealed similar seasonal patterns for the four communities (Fig. 2). Datasets from the same month clustered together in the plot, except for June datasets, thereby forming a yearly cycling of community succession. Consistent with this observation, the community dynamics were significantly correlated among the four microbial communities (Mantel test, correlation coefficients >0.75, *p* <10^-4^; Fig. S6). The average Sørensen-Dice dissimilarity between datasets from the same season ranged from 0.58 to 0.78 (Fig. 3), while that between datasets from opposite seasons (i.e. winter *vs*. summer or spring *vs*. fall) ranged from 0.77 to 0.94), indicative of greater dissimilarity of communities between opposite seasons (Fig. 3). The spring and fall communities differed from each other as much as the summer and winter communities (Fig. 3). The dissimilarities within the same season were significantly lower than those between spring and fall *(p* <10^-16^) and between summer and winter *(p* <10^-16^). However, dissimilarities between summer and winter communities did not significantly differ from those between spring and fall communities *(p* >0.1), except that spring *vs.* fall dissimilarities were higher than summer *vs*. winter dissimilarities for the prokaryotic community *(p* <10^-6^).

**Figure 2:**
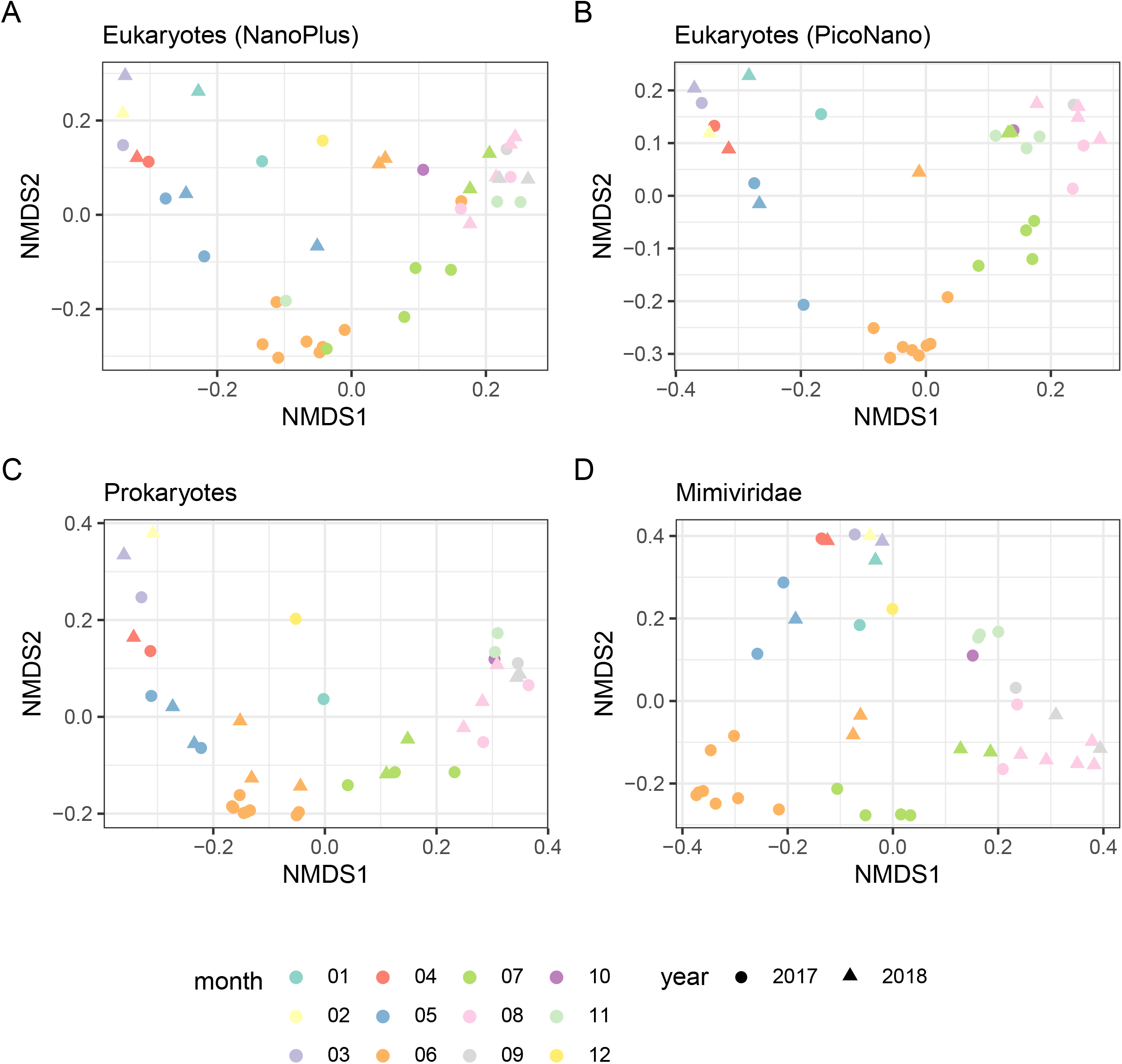
NMDS ordination of different communities. Strong seasonality for every community was detected at station “j”. Samples clustered together according to sampling months. Different colors show different months, shapes represent years. Stress of the NMDS was between 0.09 and 0.13 (prokaryotic NMDS: 0.09, PicoNano NMDS: 0.11, NanoPlus NMDS: 0.13, *Mimiviridae* NMDS: 0.11).

**Figure 3:**
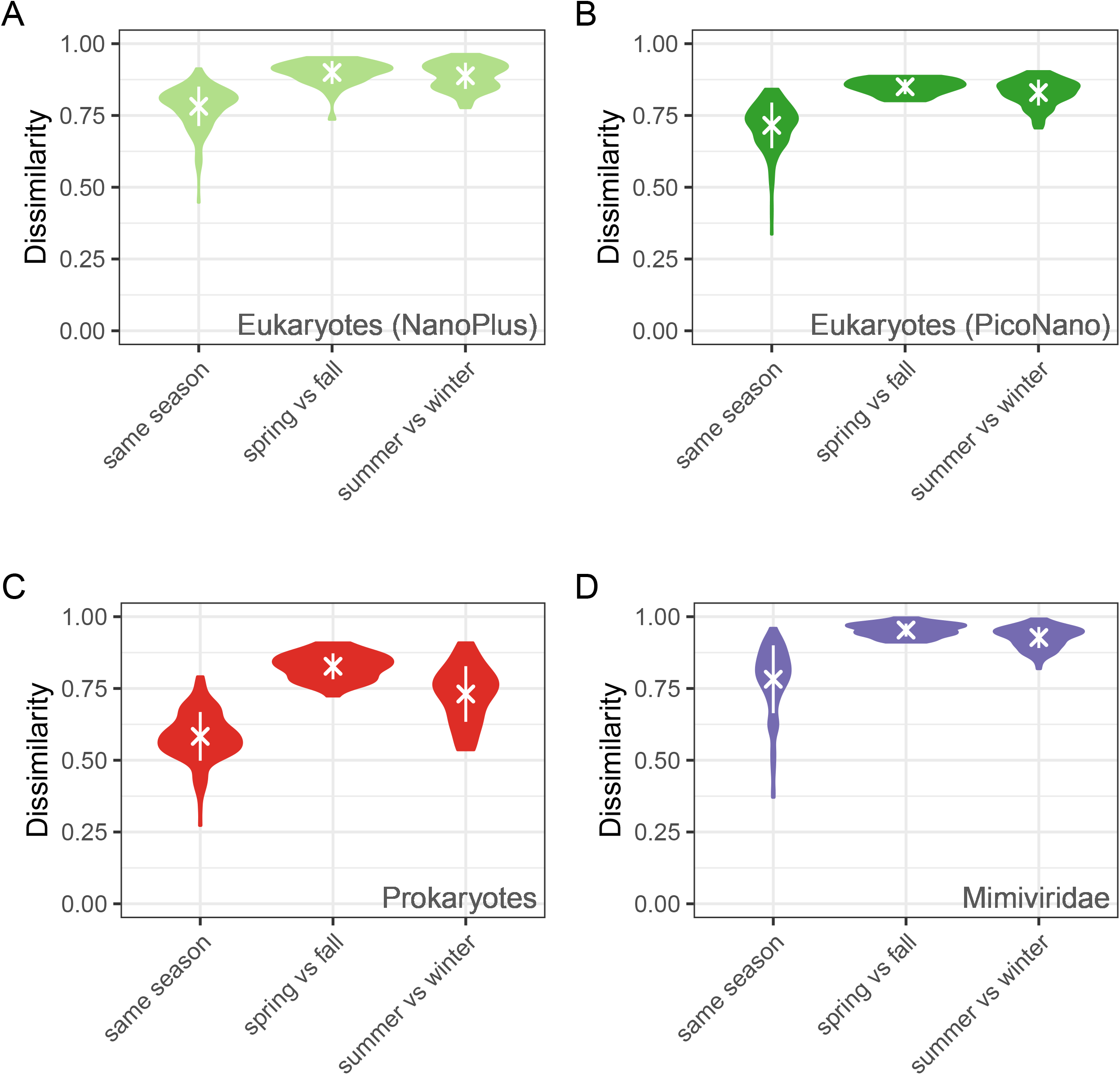
Dissimilarity metrics according to seasons. Violin plot shows dissimilarities of samples collected in opposite seasons (i.e., summer *vs.* winter or spring *vs.* fall). Communities were equally dissimilar between spring and fall as between summer and winter. Average Sørensen-Dice dissimilarity was lower between datasets from the same season (PicoNano: 0.72 ± 0.08, NanoPlus: 0.78 ± 0.06, Prokaryotes: 0.58 ± 0.09, and *Mimiviridae:* 0.78 ± 0.11; average dissimilarity ± 1 standard deviation) than between datasets from opposite seasons (PicoNano: 0.83 ± 0.04, NanoPlus: 0.89 ± 0.04, Prokaryotes: 0.77 ± 0.09, and *Mimiviridae:* 0.94 ± 0.03).

### Distinct patterns in seasonal dynamics among eukaryote, prokaryote, and *Mimiviridae* communities

The annual cycles of the communities were characterized by plotting the community dissimilarity between samples against the length of time separating the sampling events (Fig. 4A-D). In addition, we calculated the dissimilarities between communities sampled at intervals of approximately 1-month (i.e. 30 ± 10 days; hereafter D30), approximately 6 months (i.e. 182 ± 30 days; D182), and approximately one year (i.e. 365 ± 30 days; D365).

**Figure 4:**
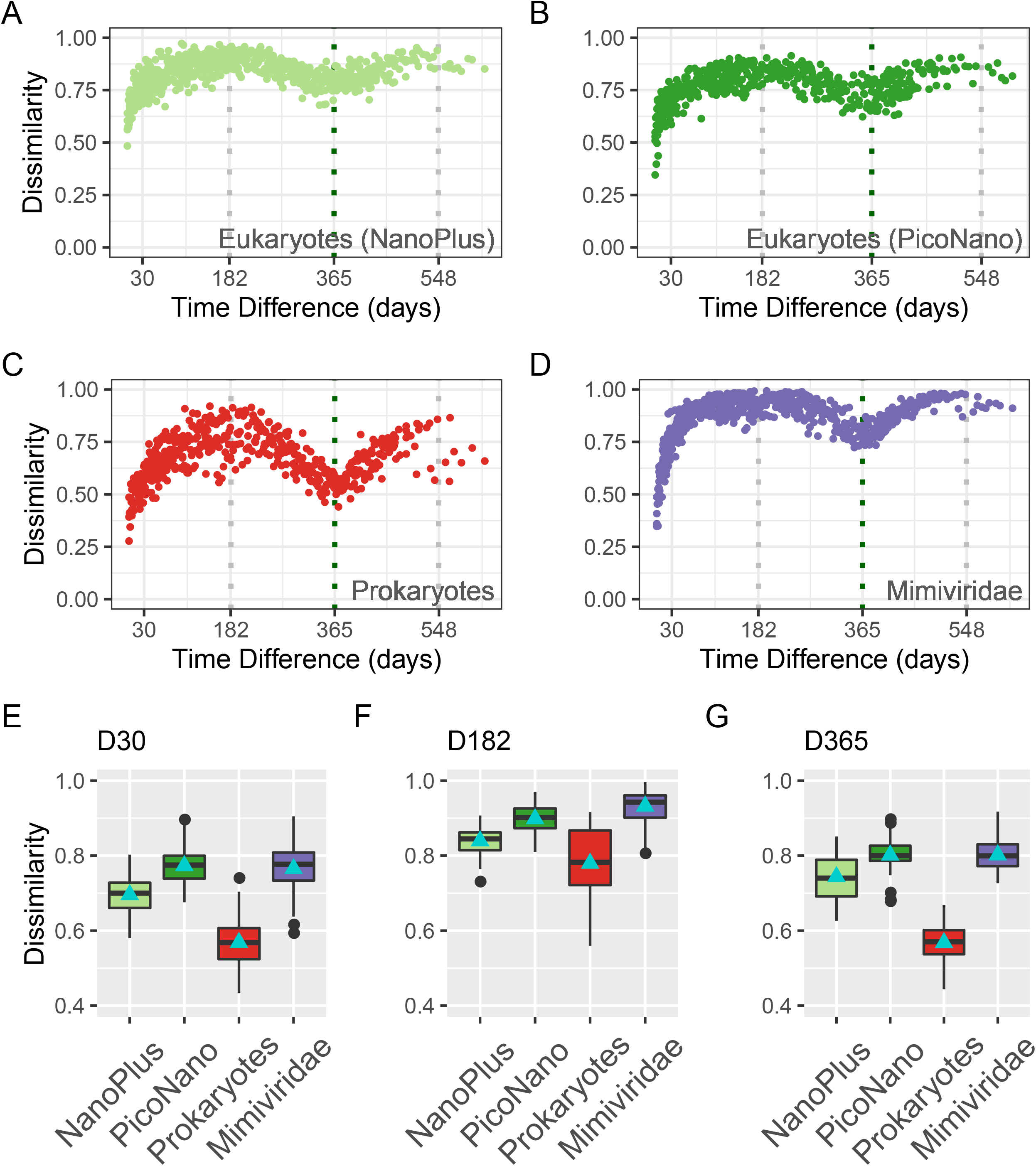
Sørensen-Dice dissimilarity metrics for the four different communities. (A-D) Dot plots showing pairwise dissimilarities of all samples plotted against sampling interval (days). (E-F) Boxplots showing dissimilarity and average dissimilarity of the four communities at three different time points.

The level of these dissimilarity measures (i.e. D30, D182, and D365) was consistent with the yearly cycle pattern observed for the four communities (Fig. 4E-G). D182 was the largest among these measures for all communities (NanoPlus, PicoNano, prokaryotes, *Mimiviridae*), and the levels of D30 and D365 were comparable. However, these measures substantially differed when compared between communities *(p* <10^-6^), except for the comparisons between *Mimiviridae* and PicoNano communities for D30 and D365. These measures were the highest for *Mimiviridae* and PicoNano communities, followed by the NanoPlus community. The prokaryotic community showed the lowest dissimilarity values for all measures (i.e. D30, D182, and D365).

### Recurrence and persistence of ASVs

To dissect the dynamics of the eukaryote, prokaryote, and *Mimiviridae* communities, we investigated temporal patterns in the occurrence of individual ASVs. First, we analyzed the temporal pattern of the four most abundant ASVs in each community (Fig. 5). This analysis revealed that the dynamics of the selected prokaryotic ASVs clearly differed from those of other ASVs. The prokaryotic ASVs were characterized by a relatively low maximum relative abundance (<31%), whilst the dynamics of the most abundant ASVs from other communities (i.e. NanoPlus, PicoNano, and *Mimiviridae)* were characterized by sharp peaks in relative abundance (up to 70%) (Fig. 5). The most abundant prokaryotic ASV was assigned to SAR11 (Clade 1a) and showed 10%–30% relative abundance more than 7 months from late autumn to early spring (present in 32 samples in total).

**Figure 5:**
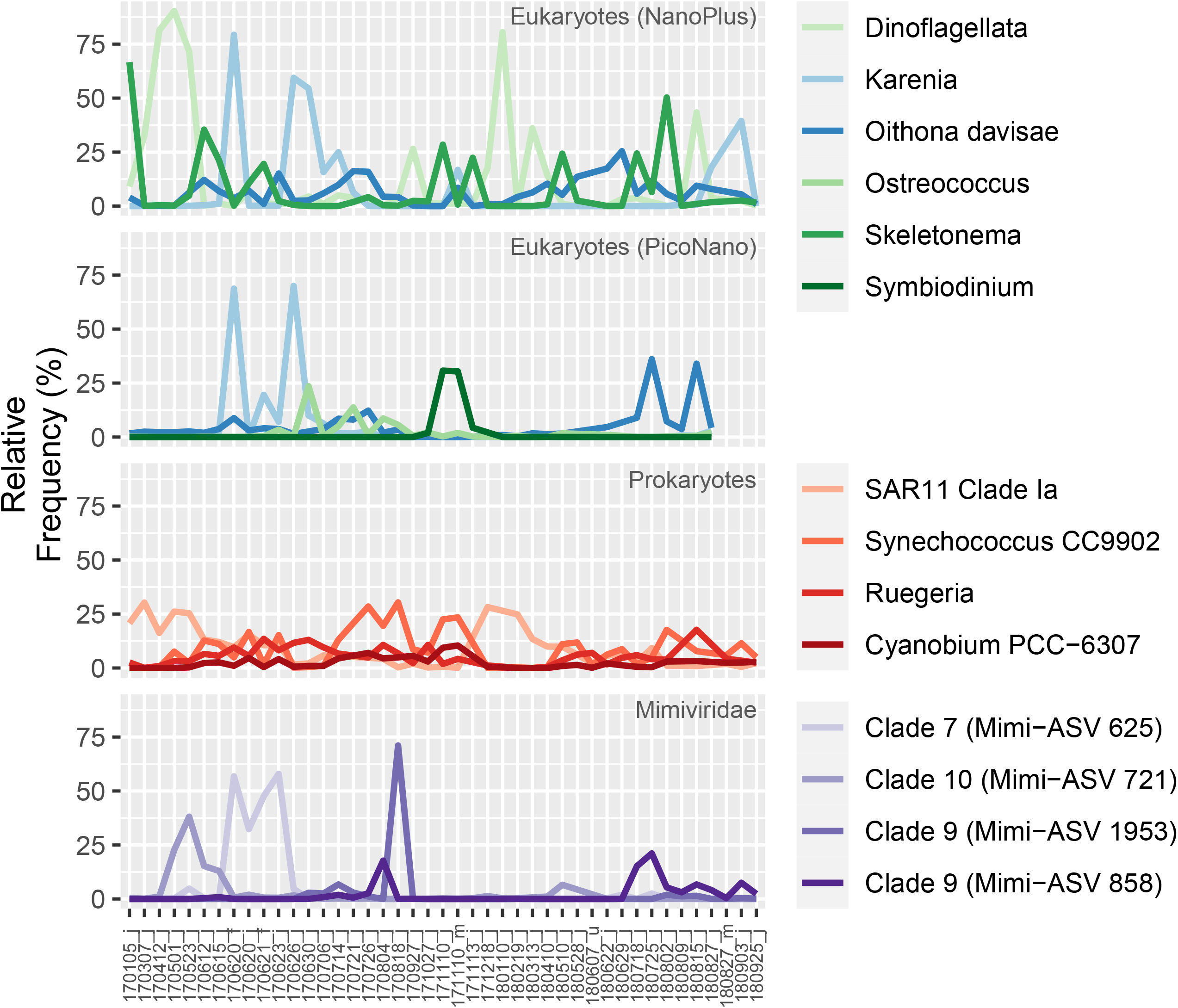
Four most abundant ASVs of the subsampled datasets of each microbe-group and their taxonomic assignment. Eukaryotes are plotted in green and blue, prokaryotes in red, and *Mimiviridae* in purple. Eukaryotes are plotted in blue if their ASV was one of the four most abundant ASVs in both size fractions.

This result prompted us to further examine recurrence (i.e. number of samples in which an ASV was detected) and persistence (i.e. maximum number of successive samples in which an ASV was detected) of ASVs. In this analysis, we considered all the ASVs with >1% relative abundance in at least one sample. Among the four microbial groups, prokaryotes showed the highest level of ASV recurrence, and *Mimiviridae* showed the lowest (Fig. 6). Similarly, prokaryotes showed longer ASV persistence compared with the other microbe groups (Fig. S7). For instance, three prokaryotic ASVs (of 73 prokaryotic ASVs) were continuously present in more than 19 datasets, while there were no such long-lasting ASVs in the *Mimiviridae* or eukaryote groups.

**Figure 6:**
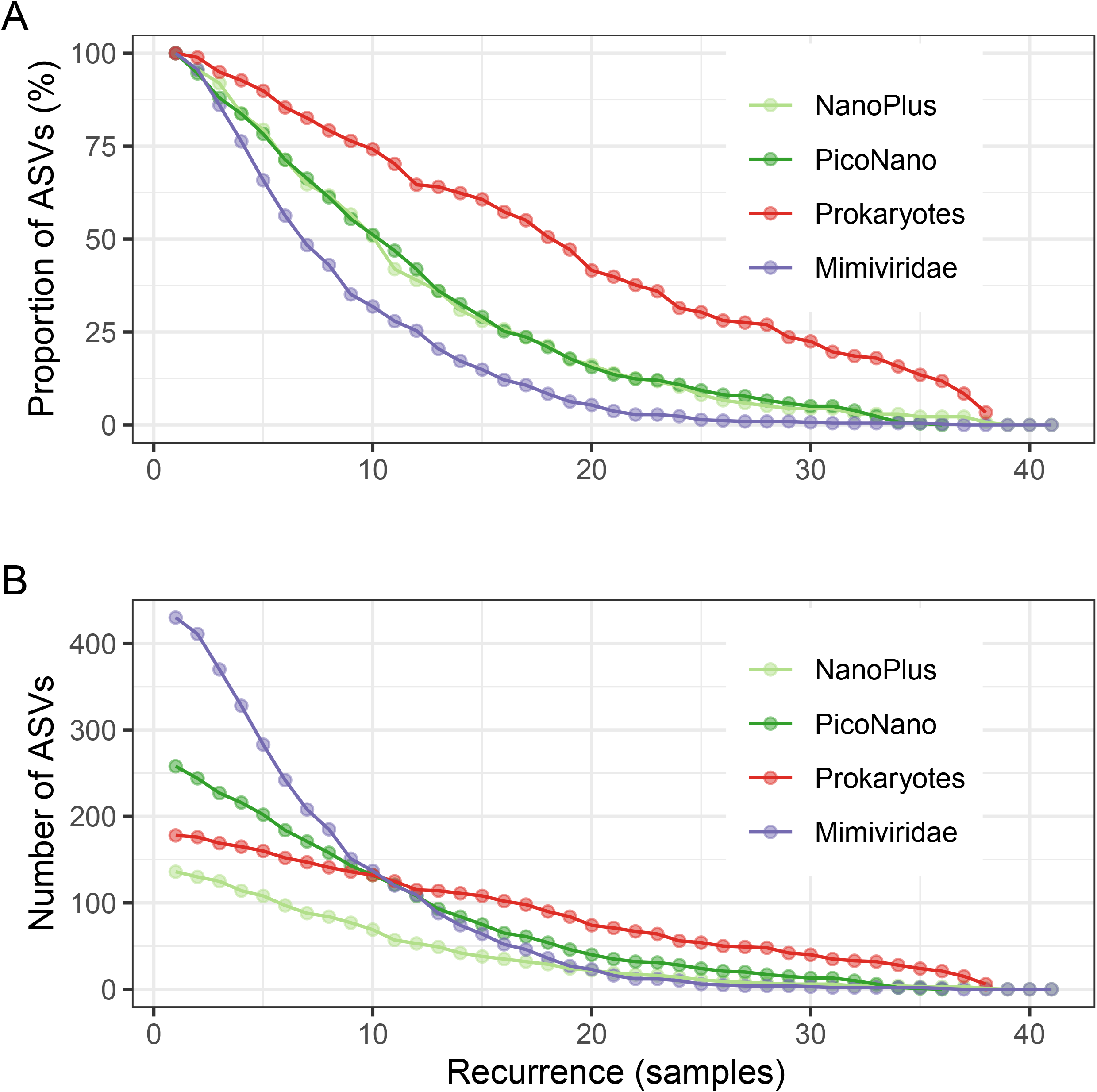
Recurrence of abundant ASVs. Line plots show number of samples in which abundant ASVs were present (i.e., recurrence). (A) Percentage of abundant ASVs and how often they were found. (B) Recurrence as indicated by absolute number of abundant ASVs and number of samples in which they were present. Most *Mimiviridae* ASVs were found in fewer than ten samples, while roughly half of all eukaryotic ASVs were found in fewer than ten samples. Most prokaryotic ASVs were found in more than half of all samples. An ASV was considered abundant if it generated at least 1% relative reads in one sample.

## Discussion

Eukaryotic communities in the study area were dominated by Metazoa, Alveolata, and Stramenopiles. These lineages are commonly found in coastal eukaryotic communities (Chen *et al*. 2017; Martin-Platero *et al*. 2018; Gran-Stadniczeñko *et al*. 2019a), although detailed compositions of microbial eukaryote communities in coastal areas vary among different locations (Chen *et al*. 2017; Martin-Platero *et al*. 2018; Gran-Stadniczeñko *et al*. 2019a). A notable feature of the study area is the occurrence of HABs (Fig. 1C), which were detected in our sequence data (Fig. 1A, B). The prokaryotic communities in the Uranouchi Inlet (Fig. 1D) were similar to those reported for other coastal areas, where the communities are mainly composed of Proteobacteria, Bacteroidetes, and Cyanobacteria (Sakami *et al*. 2016; Martin-Platero *et al*. 2018; Santi *et al*. 2019), but different from those in brackish water (Fazi *et al*. 2020) or freshwater (Bock *et al*. 2018), which are dominated by the same phyla, but different classes. Finally, the previously developed metabarcoding method (Prodinger *et al*. 2020) revealed diverse *Mimiviridae* (6261 ASVs, 464.6 ASVs per sample on average) that were grouped into 13 clades (Fig. 1E).

All four studied microbial communities, including the community of *Mimiviridae*, showed clear seasonal cycles (Fig. 2). Such seasonal changes in microbial communities have been reported previously for eukaryotes (Giner *et al*. 2019; Gran-Stadniczeñko *et al*. 2019a), bacteria (Ward *et al*. 2017; Chafee *et al*. 2018), large DNA viruses (Sandaa *et al*. 2018; Gran-Stadniczeñko *et al*. 2019b), and smaller viruses (Chow and Fuhrman 2012; Pagarete *et al*. 2013; Ignacio-Espinoza, Ahlgren and Fuhrman 2020). The greatest dissimilarity in communities in our datasets was between communities sampled at 6-month intervals (Fig. 4F), while communities were more similar when compared at approximately 12-month intervals (Fig. 4G), indicating a recovery of community structure after one year. The high dissimilarity between summer and winter datasets was expected, since physicochemical parameters such as temperature are very different between these two seasons. However, for all microbe groups, the microbial communities in spring *vs.* fall were as dissimilar as the communities in summer *vs.* winter (Fig. 3), even though the differences in water conditions (e.g. temperature, salinity, and nutrient concentrations) were less pronounced between fall and spring than between summer and winter (Fig. S1, S2). We also observed that the community compositions were relatively similar between samples collected in successive months (Fig. 2 and Fig. 4E). These observations suggest that the dynamics of microbial communities cannot be explained solely by changes in abiotic factors, but are governed by a Markovian process in which the current biotic state substantially influences the next biotic state. This pattern is obvious in macro-ecosystems; for example, a forest looks different in spring and autumn even on days with similar weather. Microbial communities with a much higher turnover rate (one to several days) appear to be no exception to this rule. A previous study on oceanic microbial communities suggested that biotic factors (such as the subset of communities) are better predictors of microbial community composition than are abiotic factors (such as geography, nutrient concentration, and temperature) (Lima-Mendez *et al*. 2015). This reinforces the classical notion that the sum of biotic and abiotic factors forms the environment that affects individual microorganisms. We suggest that the microbial communities in the Uranouchi Inlet at a given time strongly influence the formation of the next generation of communities. A similar observation and idea has been proposed for post-bloom communities (Needham and Fuhrman 2016).

All the studied microbial communities became similar to the original communities after a year, but the recovery level of community structure was far below 100% (i.e. 19% to 44% recovery, Fig. 4). In other words, more than half of the ASVs in the original communities were undetected by deep sequencing one year later. Previous long-term studies on prokaryotic (Fuhrman, Cram and Needham 2015) and viral communities (Chow and Fuhrman 2012; Ignacio-Espinoza, Ahlgren and Fuhrman 2020) detected a similar trend that continued for several years.

Therefore, it seems legitimate to postulate that the marine microbial community structure at the ASV level never perfectly recovers over time (Fig. 7A). This implies a significant role of the “invisible” rare biosphere in the formation of a new community structure through their capacity to bring forth new abundant microbes over time (Giner *et al*. 2019; Ignacio-Espinoza, Ahlgren and Fuhrman 2020). The rare biosphere may be undetectable, even with state-of-the-art metabarcoding approaches as used in this study. The technical limitations in uncovering the rare biosphere combined with its capacity to substantially replace abundant populations may explain why microbes cannot completely recover their community structure, although migration and evolution could also have effects. Interestingly, Furman et al. found that the dissimilarity of bacterial community structure increased during the first four years of analysis (Fuhrman, Cram and Needham 2015). After this period, the dissimilarity level still fluctuated (according to seasonal dynamics) but its basal level remained constant (Fuhrman, Cram and Needham 2015). This and our observations suggest the existence of a “memory” (or a historical trace) of the original community structure that influences the structures of successive communities only for a short time period (one to a few years) (Fig. 7A).

**Figure 7:**
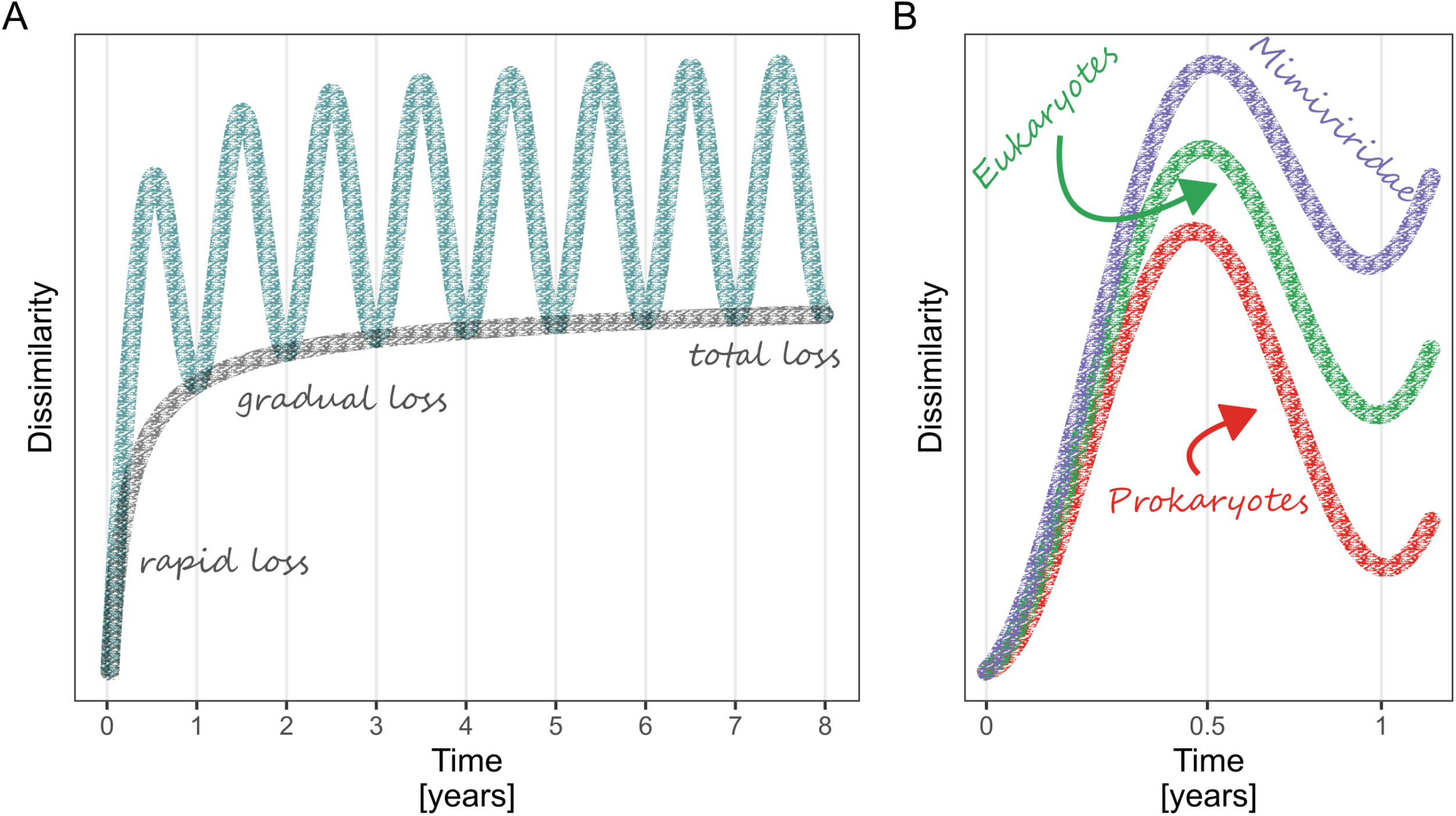
Schematic, visual representation of community memory model. (A) Seasonally dependent dissimilarity of one community over a longer sampling period with seasonal fluctuations (dark cyan) and fading community memory (grey). (B) Line plots showing changes in community dissimilarity over the short period represented in our datasets. *Mimiviridae* communities (purple) were more dissimilar than were eukaryote (green) and prokaryote (red) communities.

Although a pattern of annual cycling and partial recovery after a year was commonly observed for the four microbial communities in this study, the degree and speed of changes in the community structure showed intriguing differences among the communities (Fig. 4). A schematic diagram in Fig. 7B illustrates the conceptual and qualitative differences in the observed community dynamics among eukaryotes, prokaryotes, and *Mimiviridae* (Fig. 7B). The D182 and D365 values were highest for *Mimiviridae* and lowest for prokaryotes. Thus, in terms of “memory” of community structure, *Mimiviridae* have the shortest memory, eukaryotes have medium memory, and prokaryotes have the longest memory. The rapid change and a low level of recovery after one year for the *Mimiviridae* community contrasts with the result from a previous study on the seasonal changes of small viruses (i.e. mostly bacterial viruses), which detected strong stability of the small virus community, with 95% of viral contigs present in all samples (Ignacio-Espinoza, Ahlgren and Fuhrman 2020). In the Uranouchi Inlet, the prokaryotic community changed slowly and showed the highest level of recovery after a year. The distinct community dynamics between prokaryotes and eukaryotes in the Uranouchi Inlet is consistent with a previous study conducted on a shorter time scale (Martin-Platero *et al*. 2018); they found prokaryotic communities to be more stable and to change more slowly than eukaryotic communities.

To dissect the differences in the community dynamics, we analyzed the recurrence and persistence of ASVs among eukaryote, *Mimiviridae,* and prokaryote communities. The recurrence and persistence levels of ASVs (Fig. 6, Fig. S7) and the relative abundance of abundant ASVs (Fig. 5) differed among the microbial groups. Overall, *Mimiviridae* ASVs showed lower recurrence and persistence levels than eukaryotic ASVs. In contrast, prokaryotic ASVs showed higher recurrence and persistence levels than eukaryotic ASVs. The abundant *Mimiviridae* ASVs were found only in a few samples but they had high relative abundance (21.1%-71.3%). In contrast, the abundant prokaryotic ASVs were dominant throughout much of the year, whilst their maximum relative abundance was lower (<30%) than that of dominant ASVs in other microbial groups (Fig. 5). The pattern of eukaryotic ASVs occurrence was between these extremes. The four most abundant ASVs in the PicoNano and the NanoPlus groups had high relative abundance (23.5%-90.3%) but were present in more samples than were the most abundant *Mimiviridae* ASVs (Fig. 5). Therefore, the systematic differences in the recurrence/persistence levels of individual ASVs are consistent with the differences in the dynamics at the community level.

The differences in the community and ASV dynamics among eukaryotes, *Mimiviridae,* and prokaryotes may result from the limitations of metabarcoding. It is worth noting that ASVs may have different resolutions across microbial groups. For instance, the result that bacterial ASVs showed longer persistence than eukaryote and *Mimiviridae* ASVs may because their resolution was lower. However, if we assume that this is the case, the comparatively high abundance peaks for the major eukaryotic and *Mimiviridae* ASVs become difficult to compromise, because their resolution was assumed to be higher than that of bacterial ASVs. Thus, the differences in the ASV community dynamics likely reflect some essential features of the dynamics of the populations belonging to the respective microbial communities. In the following paragraph, we attempt to explain these differences in light of the possible ecological strategies of individual populations within those communities.

The population dynamics of microbes largely depend on resource availability and their survival strategies, which have historically been classified into “*r*” and “*K*” strategies. The original interpretation of *r/K* selection theory states that organisms maximize their fitness through either high reproduction rates (high *r*), or the ability to constantly exist in relatively high numbers (high carrying capacity, *K*) (Andrews and Harris 1986). In microbial ecology, the *r*/*K* nomenclature has often been used more qualitatively (Andrews and Harris 1986). That is, “*r*-strategists” are specialized to rapidly consume resources and reproduce quickly (Andrews and Harris 1986) so that they thrive through an influx of nutrients (Santi *et al*. 2019). In contrast, *K*-strategists adapt to a stable environment with limited resources (Andrews and Harris 1986). They tend to show slow growth rates and their population dynamics are more stable than those of *r-* strategists (Endo, Ogata and Suzuki 2018).

In our study, several of the major eukaryotic ASVs were present over a long period of time (i.e. the *Skeletonema* sp. ASV, the *Oithona davisae* ASV and the Dinoflagellata ASV; Fig. 5), reminiscent of *K*-strategists. The Dinoflagellata ASV in the NanoPlus fraction was abundant throughout several months in spring during oligotrophic conditions. The *Oithona davisae* ASV (a copepod preying on plankton) was present throughout the year with no pronounced peak in its relative abundance. In contrast, other abundant ASVs (e.g. the Symbiodinium ASV) and those assigned to HAB species (e.g. the *Chattonella* spp. ASVs, the *K. mikimotoi* ASVs, and the *H. akashiwo* ASVs; Fig. 1A and B) emerged only occasionally but with a high relative abundance (>20%). The dynamics of the HAB ASVs were consistent with their cell counts (Fig. 1C). They formed blooms only during the months with high nutrient availability (Fig. S1, Fig. S2). This ecological characteristic is typical of *r*-strategists (Andrews and Harris 1986; Endo, Ogata and Suzuki 2018).

The majority of *Mimiviridae* ASVs showed low persistence and occasional high peaks in relative abundance. The most abundant *Mimiviridae* ASVs occurred only in a few samples but had high relative abundance of over 70% (Fig. 5). These dynamics are reminiscent of *r*-strategists that show a rapid boom and bust appearance. The lower recurrence and persistence of *Mimiviridae* ASVs than eukaryote ASVs likely originate in the fact that the presence of viruses depends on the presence of their hosts, not vice versa. Furthermore, the possible existence of multiple *Mimiviridae* that compete for the same eukaryotic host species (Baudoux and Brussaard 2005) may lead to the *r*-strategist like feature in the dynamics of *Mimiviridae* ASVs.

A recent study suggested that *Mimiviridae* that infect *K*-strategist eukaryotes have long infection and latent periods to maximize the chance of vertical transmission, since the chances of horizontal transmissions are low due to the low host density (Blanc-Mathieu *et al*. 2020). Viruses of *K*-strategist eukaryotes are more likely to exist inside the host cell (i.e. a long latent period), while viruses of *r*-strategist eukaryotes are more likely to be found outside their host (i.e. they show higher virulence and tend to produce virions more quickly) (Blanc-Mathieu *et al*. 2020). The size fraction we used for the detection of viral sequences (0.2-0.8 μm) may be enriched in free virions, which might have led to the preferential detection of *r*-strategist *Mimiviridae.* A few *Mimiviridae* ASVs showed a relatively long persistence (occurrence in more than ten continuous samples) (Fig. S6) or a high recurrence level (Fig. 6). Investigation of active *Mimiviridae* (e.g. by RNA-Seq from the PicoNano and NanoPlus fractions) may reveal *Mimiviridae* ASVs showing *K*-strategist dynamics.

The most abundant prokaryotic ASVs had comparatively low relative abundances (Fig. 5), but were found in many samples (Fig. 6). The most abundant prokaryotic ASV belonged to SAR11, which is a *K*-strategist characterized by its ubiquitous presence in the ocean and low maximum growth rate (Suttle 2007; Giovannoni 2017). As expected, this ASV was abundant during months with low nutrient availability with no pronounced sharp peaks in abundance. Several bacterial genera are known to be *r-*strategists (e.g. *Ulvibacter, Formosa,* and *Polaribacter*) (Teeling *et al*. 2012, 2016; Needham and Fuhrman 2016), but they were not highly represented in our dataset. In this study, prokaryotic communities were more stable than were eukaryotic and *Mimiviridae* communities. Heterotrophic prokaryotes depend on organic substrates especially from eukaryotes, but their interactions are mostly indirect and less specific compared with the specificity of *Mimiviridae-eukaryote* interactions. Overall, the distinct community dynamics among eukaryotes, *Mimiviridae*, and prokaryotes revealed in our study may be due to the differences in specificity of interactions as well as the prevalence of *r vs. K* strategies among these microbial groups.

In summary, our study revealed synchronic seasonal cycles for eukaryote, *Mimiviridae*, and prokaryote communities in the Uranouchi Inlet. The seasonality could not be explained solely by abiotic factors and is suggested to be shaped through a Markovian process of successive communities under the influence of abiotic factors. Despite their synchronized community dynamics, we discovered intriguing differences in the dynamics in several aspects, including the length of “community memory” among eukaryotes, *Mimiviridae,* and prokaryotes. These differences at the community level were consistent with the recurrence and persistence of individual ASVs in the respective communities. Therefore, the distinct community dynamics among eukaryotes, *Mimiviridae*, and prokaryotes likely originate in systematic differences in the specificity of interactions (i.e. virus*-*eukaryote *vs* prokaryote*-*eukaryote) as well as the dominant survival strategies across the studied microbial groups. Overall, our results provide an important step towards mechanistically understanding the community dynamics of microorganisms in light of ecological strategies of individual microorganisms.

## Acknowledgments

F.P. performed experiments, bioinformatics analyses, and wrote the initial manuscript. H.E., K.N., T.Y. and H.O. designed the work, helped to interpret the results, and improved the manuscript. Y.G. and H.T. contributed to sequencing. R.B.M and Y.L. contributed to bioinformatics analysis. K.T. and I.T. contributed to experimental analyses. Y.T., K.N., and E.T. contributed to sampling. All authors contributed to writing and agreed to the final manuscript.

## Funding

This work was supported by The Canon Foundation [No. 203143100025]; JSPS/KAKENHI [Nos. 26430184, 17H03850, 18H02279, and 16H06279 (PAGS)]; Scientific Research on Innovative Areas from the Ministry of Education, Culture, Science, Sports and Technology (MEXT) of Japan [Nos. 16H06429, 16K21723, and 16H06437]; The Kyoto University Foundation, and the Collaborative Research Program of the Institute for Chemical Research, Kyoto University [Nos. 2019-33, 2018-31, 2017-25, and 2016-28]. Computational work was completed at the SuperComputer System, Institute for Chemical Research, Kyoto University. We thank the Kochi Prefectural Fisheries Research Institute for providing environmental metadata. We thank Jennifer Smith, PhD, from Edanz Group (https://en-author-services.edanz.com/), for editing a draft of this manuscript.

The authors declare no conflict of interest.

## Data availability

The raw sequencing data of this study have been deposited in the DNA database of Japan (accession number DRA010976). The environmental data and cell count data are available in the Microsoft Excel file (dataset S1). The amplicon data analysis pipelines for eukaryotes, prokaryotes, and *Mimiviridae* have been deposited online and are available for download (www.github.com/FlorianProdinger/pipeline_18S, www.github.com/FlorianProdinger/pipeline_16S, www.github.com/FlorianProdinger/MAPS2, 17 February 2021, date last accessed).

---

Fig. S1: Cell counts, biotic, and abiotic data collected over 6 years. Blue dotted lines indicate sampling period during which amplicon sequencing was performed.

Fig. S2: Monthly and seasonal summary of cell counts, biotic, and abiotic data. (A-L) Boxplots showing monthly and seasonal biotic and abiotic data. (M) Bar plot showing monthly sum of daily cell counts throughout 6 years.

Fig. S3: Rarefaction curves of all samples. (A) Rarefaction curves for subsampled datasets. (B) Rarefaction curves for data without subsampling.

Fig. S4: Visualization of different diversity indices across all datasets subsampled at 8000 reads. (A) Shannon’s diversity index; (B) Richness of different communities; (C) Pielou’s evenness index.

Fig. S5: Phylogenetic tree with *Mimiviridae* ASV placements. Names of leaves correspond to known reference *Mimiviridae*. Unnamed leaves are PolB amino acid sequences assembled from metagenomic data. Red dots indicate placement of sequences generated from *Mimiviridae PolB* amplicon sequencing.

Fig. S6: Correlations of dissimilarity between different communities. All communities are statistically significantly correlated with each other *(p* <10^-4^/

Fig. S7: Persistence of abundant ASVs in different microbial communities. (A) Percentage of abundant ASVs that were persistent (i.e., continually found in a series of samples). (B) Absolute number of abundant ASVs and their persistence.

Dataset S1: Microsoft Excel file contains environmental data and cell count data obtained from the Kochi Prefectural Fisheries Research Institute.

## Notes

### Competing Interest Statement

The authors have declared no competing interest.

### Summary of Updates

We changed the format according to our targeted journal. We revised some sentences and paragraphs to be clearer and provide a better explanation of our findings. The main massage or figures were not changed.

