## Supplementary figures and images for "Year-round dynamics of amplicon sequence variant communities differ among eukaryotes, *Mimiviridae*, and prokaryotes in a coastal ecosystem"

### Fig. S1

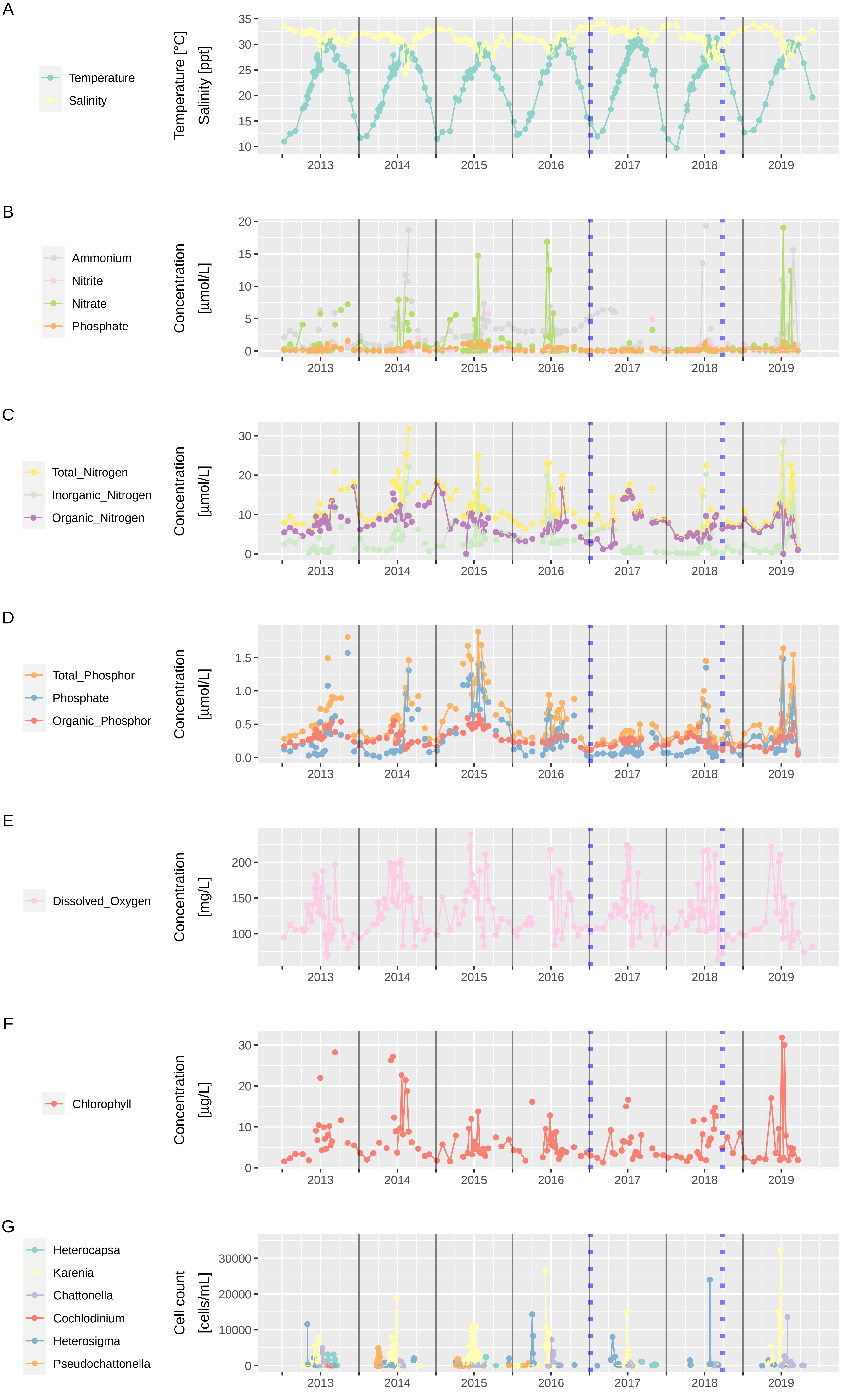

### Fig. S2

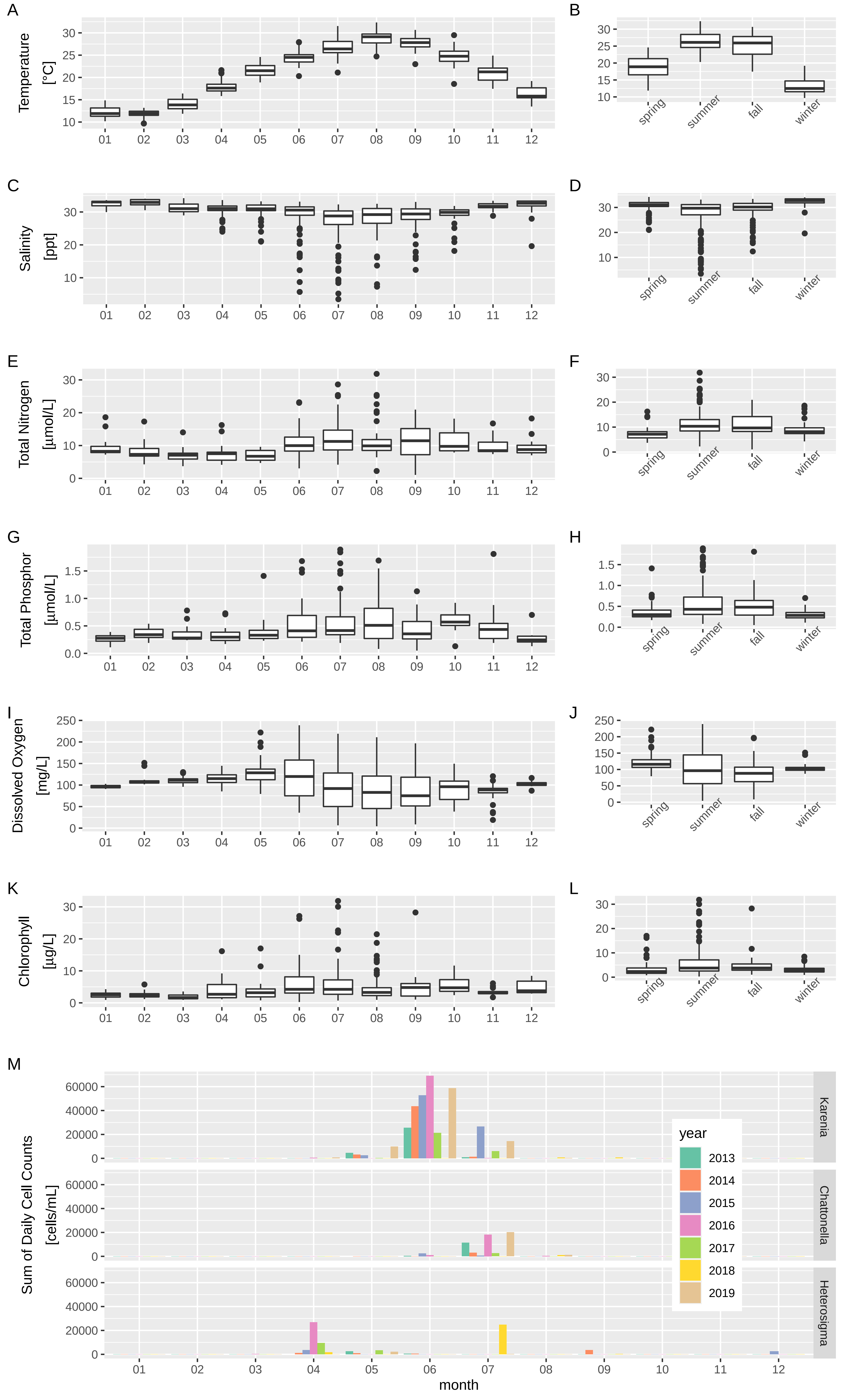

### Fig. S3

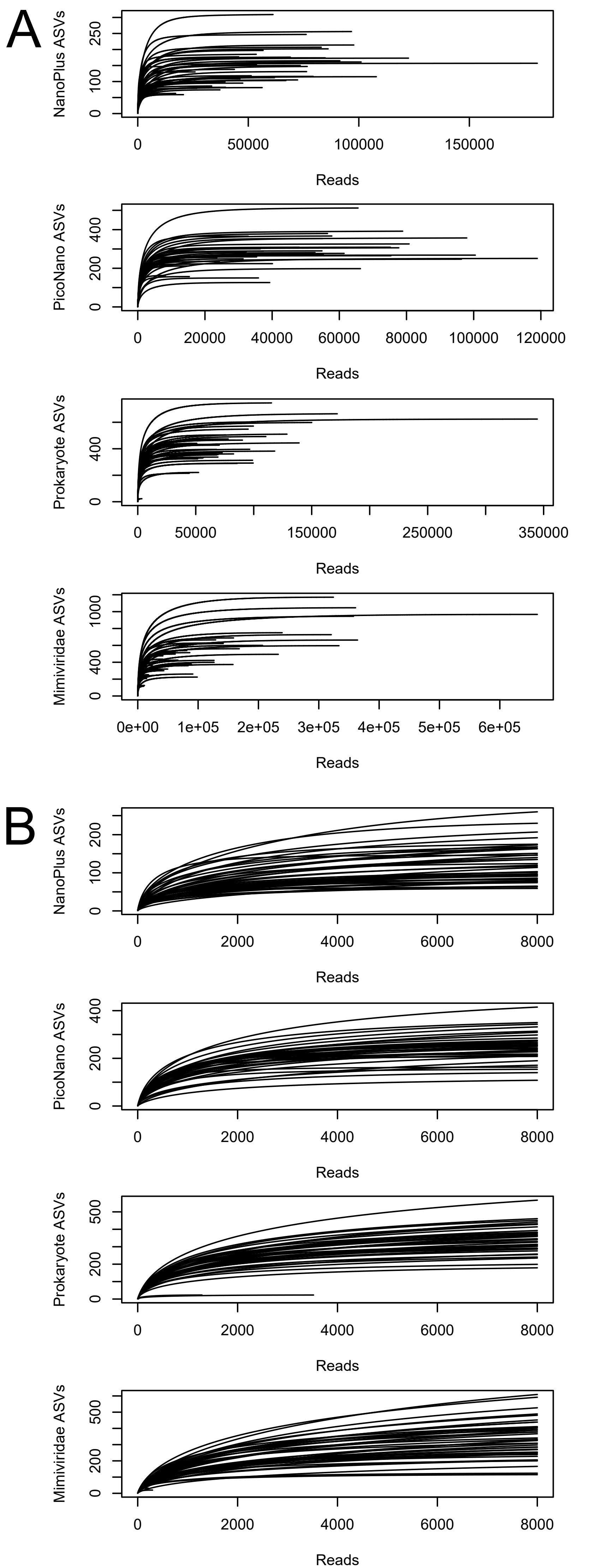

### Fig. S4

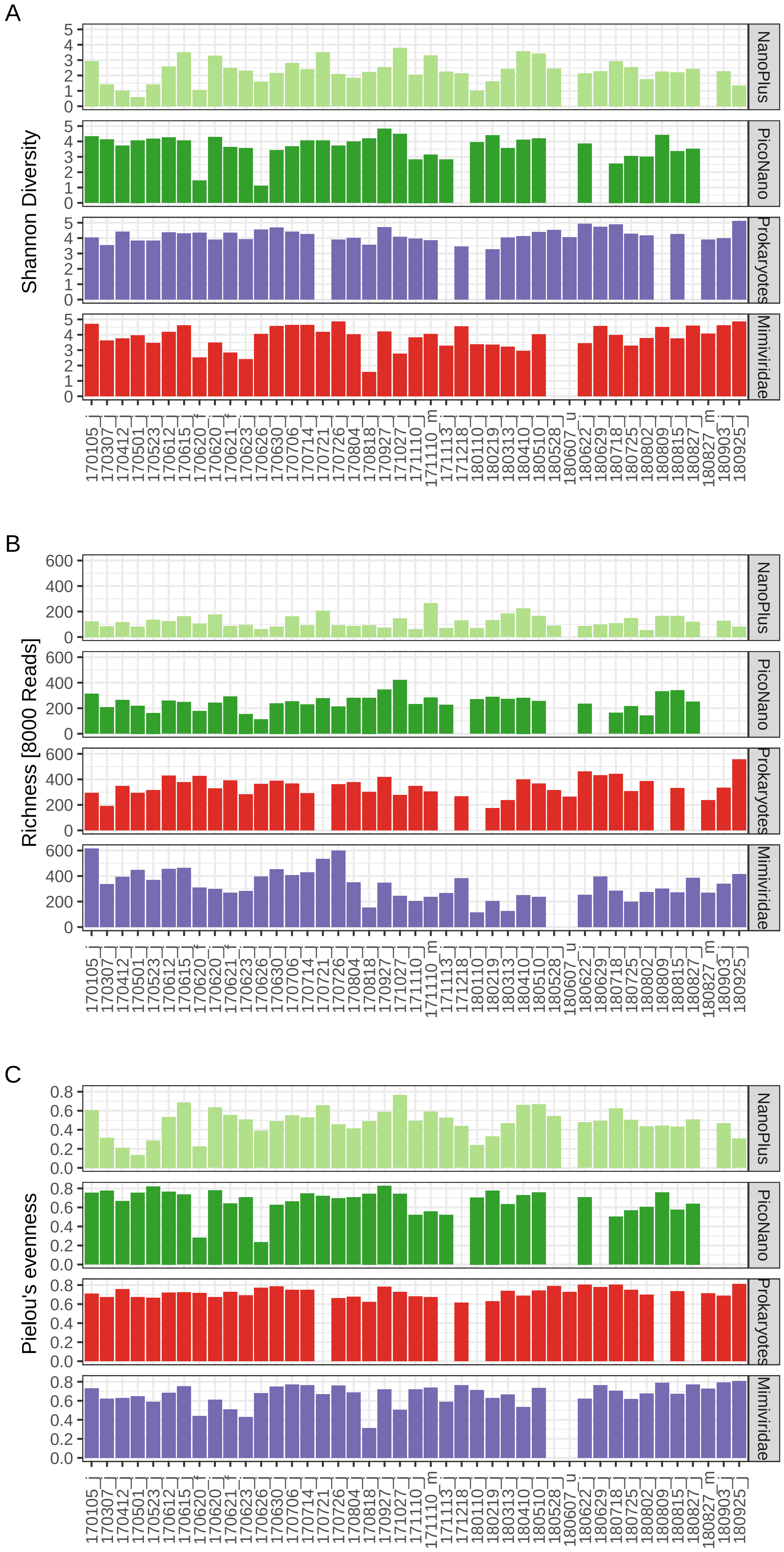

### Fig. S5

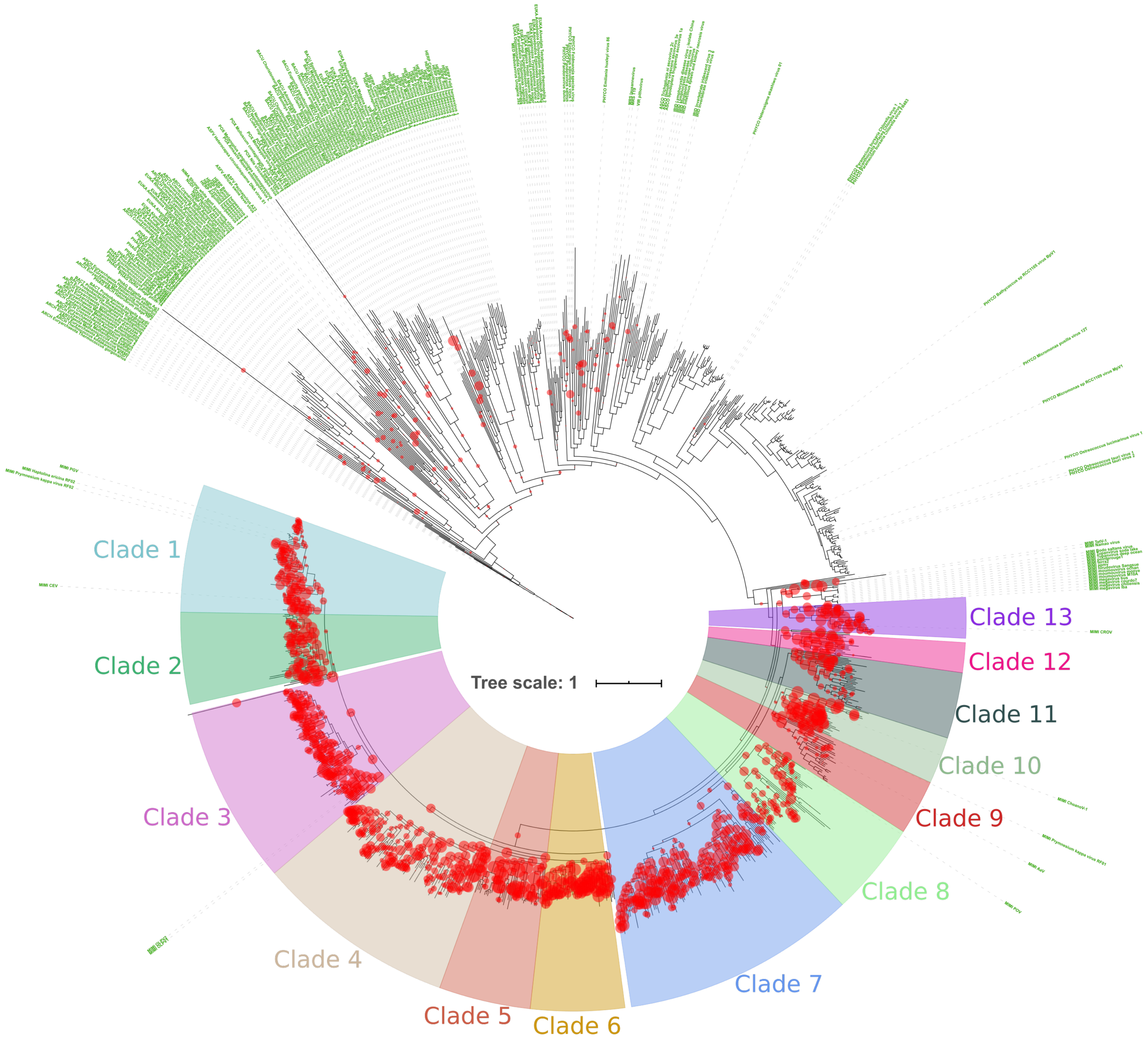

### Fig. S6

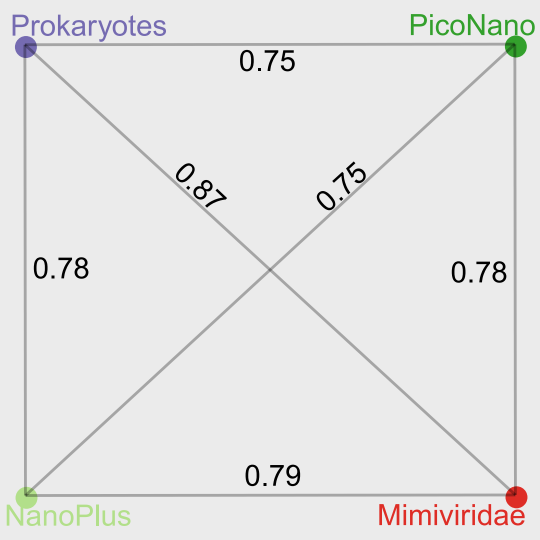

### Fig. S7

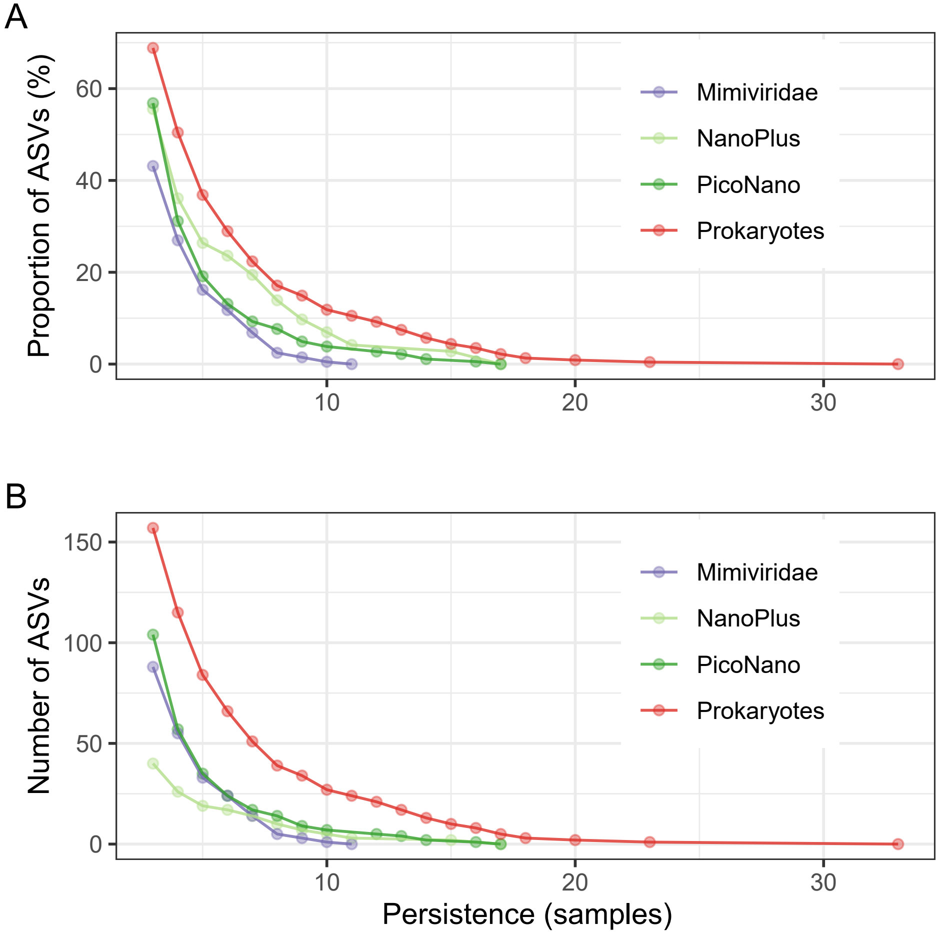
